# Rare species do not disproportionately contribute to phylogenetic diversity in a subalpine plant community

**DOI:** 10.1101/2024.11.01.621409

**Authors:** Leah N Veldhuisen, Verónica Zepeda, Brian J Enquist, Katrina M Dlugosch

## Abstract

**Premise of the study:** Worldwide, 36% of all plant species are exceedingly rare, and are at the highest risk of extinction. Extinction risk has also been observed to be phylogenetically clustered, threatening the loss of lineages that contribute unique evolutionary diversity to communities. Rare plants’ contributions to phylogenetic diversity are largely unknown, however. We investigate whether rare species (as measured by local abundance and range size) contribute disproportionately to phylogenetic diversity in a subalpine plant community.

**Methods:** We collected abundance data at three sites in Washington Gulch near the Rocky Mountain Biological Laboratory (RMBL, Gothic, Colorado, USA) in 2021 and 2022. We then calculated range size for each species. We calculated phylogenetic signal for abundance and range size, compared community phylogenetic metrics weighted by range size and abundance to unweighted metrics, and quantified the change in phylogenetic diversity when removing single species and groups of species ranked by rarity.

**Key results:** We found phylogenetic signal for abundance, but not range size. There was no difference between rarity-weighted and unweighted phylogenetic diversity metrics. Finally, phylogenetic diversity did not decline more when we removed single rare species and groups of rare species than when we removed single common species and groups of common species.

**Conclusions:** Overall, we found that rare species do not disproportionately contribute to phylogenetic diversity. This suggests that rare species likely provide phylogenetic redundancy with common species, and losing rare species may not cause disproportionate drops in phylogenetic diversity in this system.

## INTRODUCTION

Worldwide, 36% of all plant species are exceedingly rare (observed only five or fewer times globally; Enquist et al., 2019), and are at the highest risk of extinction (Yessoufou and Davies, 2016; Enquist et al., 2019). At the same time, extinction risk has been observed to be phylogenetically clustered (Willis et al., 2008; Eiserhardt et al., 2015; González-Orozco et al., 2016), threatening the loss of lineages that contribute unique evolutionary diversity to communities. While rare species can perform critical ecosystem functions, such as supporting pollinators, enhancing fire resilience, and contributing to food web diversity (Bracken and Low, 2012; Mouillot et al., 2013; Leitão et al., 2016), the full extent of their exact ecological contributions is unclear (Violle et al., 2017; Dee et al., 2019). Further, their role in maintaining phylogenetic diversity remains poorly understood, with few studies addressing this relationship (see Anderson et al., 2004; Mi et al., 2012; Herzog and Latvis, 2022).

Understanding phylogenetic diversity is essential because it is strongly linked to ecosystem processes. It correlates more positively with biomass than species richness or functional diversity (Cadotte et al., 2008), and promotes ecosystem stability, productivity, and resistance to invasion (Cadotte et al. 2012; Davies et al. 2016; Cadotte 2013; Galland et al. 2019; Delavaux et al. 2023). In plant communities specifically, phylogenetic diversity can increase diversity in other taxonomic groups (Dinnage et al., 2012; Milcu et al., 2013). Phylogenetic diversity also reflects niche differences and captures variation missed with single-trait approaches (E-Vojtkó et al. 2023; Cadotte et al. 2009; Huang et al. 2020). Given its ecological importance, phylogenetic diversity has become a key focus for conservation efforts (Tucker et al., 2012; Molina-Venegas et al., 2021). Despite the increasing use of phylogenetic diversity, it is unclear whether rare species disproportionately contribute to phylogenetic diversity, especially in the face of ongoing species losses (White et al., 2023). This study addresses this gap by investigating the role of rare species in contributing to phylogenetic diversity in a subalpine plant community via two major axes of rarity: local abundance and range size.

Rarity can be categorized in multiple ways, including small population size, limited range size, narrow habitat specificity, or combinations thereof (Rabinowitz, 1981). These metrics can independently or jointly categorize species as rare, but they do not always correlate (Sexton et al., 2009). It is unclear whether rare species ranked by range size would impact phylogenetic diversity in the same way as rare species ranked by abundance. For example, a species may have a very high local abundance but a very restricted distribution or vice versa. Species’ range sizes are dynamic; ranges can shift, shrink, or expand with global change (Hardie and Hutchings, 2010; Chen et al., 2011; Sheth and Angert, 2018). Like low-abundance species, species with small ranges are also at higher risk of extinction (Harnik et al., 2012; Manes et al., 2021). Despite this, the contribution of range-restricted species to phylogenetic diversity has not been systematically tested.

Rare species may contribute disproportionately to phylogenetic diversity. Low abundance and small-ranged species may be more phylogenetically diverse than random (overdispersed) in relation to each other and their larger community (Mi et al., 2012). The niche differentiation hypothesis suggests that each species within a community occupies its own niche, that rare species use distinct resources, and that rare species will be distinct from each other and more abundant species (Hanski, 1982; Gaston, 1994; Grime, 1998). However, recent studies show that phylogenetic diversity does not consistently serve as a proxy for functional diversity (Gerhold et al., 2015; Cadotte et al., 2019; E-Vojtkó et al., 2023). Nevertheless, phylogenetic diversity can capture community-level functional differences and link to ecosystem properties (Cadotte et al. 2009; E-Vojtkó et al. 2023; Cadotte et al. 2008; Davies et al. 2016). Testing whether rare species enhance phylogenetic diversity is essential for understanding their role in ecosystems, particularly in the context of global change.

Previous studies suggest rare species can be ecologically or functionally distinct, but their phylogenetic contributions remain understudied. Jiao et al. (2017) found that rare microbes have higher alpha and beta diversity than their abundant counterparts, though the authors did not examine phylogenetic diversity. Similarly, Ge et al. (2019) found that in desert annual plants, rare species do not closely exhibit a well-established functional tradeoff between water use efficiency and relative growth rate seen in the dominant species, suggesting niche differentiation in physiological traits (Ge et al., 2019). Like Jiao et al. (2017), Ge and colleagues (2019) did not test the relatedness of species evading the tradeoff, leaving open an important question about how phylogenetic relatedness may explain differences in rare species’ ecological strategies. In tropical tree plots worldwide, rare species were phylogenetically more diverse than abundant species, although this only held true in less disturbed plots (Mi et al., 2012). Anderson et al. (2004) showed that abundance correlated with phylogenetic relatedness in yeast communities in decaying cactus. Still, these patterns varied depending on which subfamily of cactus the yeast was growing within (Anderson et al., 2004).

In contrast, rare species may be phylogenetically redundant, randomly distributed across the phylogeny, and not disproportionately contribute to overall phylogenetic diversity. In this case, we would not expect low abundance and small-ranged species to disproportionately contribute to overall phylogenetic diversity. Instead, they would provide phylogenetic redundancy with more abundant species. In tall grass prairies, rare species did not contribute disproportionate phylogenetic diversity, providing phylogenetic redundancy within the community (Herzog and Latvis, 2022). These contrasting findings underscore the need for more focused studies on the role of rare species in contributing to phylogenetic diversity.

This study tests whether rare plant species, defined by low abundance and small range size, disproportionately contribute to phylogenetic diversity in subalpine plant communities. We hypothesize that rare species contribute more to phylogenetic diversity due to their evolutionary distinctiveness and predict that removing rare species will cause greater declines in diversity metrics compared to common species. We test this idea across an elevational gradient in the Colorado Rocky Mountains near the Rocky Mountain Biological Laboratory (RMBL, Gothic, Colorado, USA). We first predict (1) that range size and local abundance will show phylogenetic signals, indicating that common and rare species occupy different phylogenetic lineages. Then, we predict (2) that weighting phylogenetic diversity metrics by high abundance or large range size will decrease phylogenetic diversity, consistent with the prediction that rare species contribute more to diversity than common species. Finally, we predict (3) that removing species according to rarity will decrease phylogenetic diversity more when the removed species are rare than when they are common.

## MATERIALS & METHODS

### Study system & abundance data

We collected abundance data for an angiosperm community at three sites in Washington Gulch near the Rocky Mountain Biological Laboratory (RMBL, Gothic, Colorado, USA) from June to August in 2021 and 2022. RMBL is located in the East River valley of the West Elk mountains, approximately 10 kilometers from Crested Butte, Colorado. Study sites were located at 2815 m (38°53’50“N, 106°58’43“W), 3165 m (38°57’38“N, 107°01’53“W), and 3380 m (38°58’10“N, 107°01’53“W) in elevation. At each of the three sites, we had five 1.2 m x 1.2 m plots that were randomly placed within each site each summer (see more detailed description of sites and plots in Veldhuisen et al., 2023 and Sloat et al., 2015).

We identified all vascular plants to species level within each 1.2 m x 1.2 m plot at the three sites. We confirmed our identifications against RMBL Herbarium specimens (herbarium code RMBL) and iNaturalist observations. Each plot was sampled once per year near the peak of the growing season (approximately mid-July, depending on the year and elevation). Plant sampling dates were staggered to ensure all communities had the same relative phenological time point. In each plot, we counted all individuals of every species. We only counted plants rooted within the plot, and unknown and non-native species were excluded from all analyses. To quantify abundance, we averaged local abundance across all five plots at each site and the two data collection years. We did not include invasive species or species only identified to genus in our analyses.

### Range size

We calculated range size as Extent of Occurrence (EOO) and Area of Occupancy (AOO) (Gaston and Fuller, 2009; Staude et al., 2020). To obtain these values, we downloaded occurrence data for each species using the ‘RGBIF’ package (Appendix S1; Chamberlain and Boettiger, 2017) and calculated AOO and EOO with the ‘red’ package (Cardoso, 2017). We restricted GBIF downloads to occurrences in North America with <1000m coordinate uncertainty and observations made since 1990 to ensure coordinate accuracy. We used only the AOO values to calculate phylogenetic signal (Prediction 1) and rank species for testing the impact of their removal (Prediction 2), because AOO is less sensitive to extremes and outlier observations (Gaston and Fuller, 2009; Staude et al., 2020).

### Phylogenetic metrics

To calculate phylogenetic signal and community phylogenetic diversity, we used the ‘ALLMB.tre’ angiosperm phylogeny from Smith and Brown (2018). This phylogeny includes all genera in our dataset and 54 of 68 species. If a species was missing from the phylogeny but was the only member of its genus in the plots, we replaced it with a closely related congener in the Smith and Brown phylogeny (Appendix S2). Only the genus *Senecio* had multiple species in the plots missing from the phylogeny. We replaced *Senecio integerrimus* Nutt. with sister species *S. triangularis* Hook (Pelser et al., 2007). We removed *S. crassulus* A. Gray and *S. bigelovii* A. Gray from all phylogenetic analyses because they have no available genetic or phylogenetic information. We could not find an up-to-date genus-level phylogeny for *Erigeron* to inform replacements, so we removed *Erigeron elatior* and *Erigeon glacialis* from all analyses.

We calculated phylogenetic signal for both abundance and range size with Blomberg’s K (Blomberg et al., 2003) and Pagel’s λ (Pagel, 1999). We calculated Blomberg’s K with the “phylosignal()” function in the ‘picante’ R package (Kembel et al., 2010) with 5000 randomizations, and Pagel’s λ with the function “phylosig()” in the ‘phytools’ R package (Revell, 2011) with 5000 iterations and model set to “lambda.” Blomberg’s K or Pagel’s λ values near 1 indicate trait evolution consistent with Brownian motion. In contrast, K less than 1 or Pagel’s λ near zero suggests less phylogenetic signal than expected by Brownian motion.

We calculated Faith’s phylogenetic diversity (PD; (Faith, 1992), mean phylogenetic distance (MPD), and mean nearest taxon distance (MNTD) and their associated standard effect sizes (SES) in ‘picante’ (Kembel et al., 2010). Faith’s Phylogenetic Diversity (Faith, 1992) is the total evolutionary distance (branch length) across a phylogenetic tree describing the relationships among the taxa in a dataset. This metric will increase if relationships are more distant across the tree (species are more evolutionarily divergent). Mean Phylogenetic Distance is the mean distance between pairs of species. As such, it also increases as evolutionary relationships are more distant across the tree. However, it will be less sensitive to rare cases of distantly-related species that would otherwise elevate Faith’s Phylogenetic Diversity. Mean Nearest Taxon Distance is the mean of the distance to the closest relative of each taxon, and reflects the typical distance to any species’ closest relative in the dataset and increases as closest relatives are more distant. Therefore, MNTD is more reflective of the tendency of species to have closer relatives than it is of the overall evolutionary diversity across the entire tree (community of species) that is captured by the former metrics (for further discussion, see review by Tucker et al., 2017).

To calculate SES, we used each metric’s “sample.pool” null model to randomly draw species from the RMBL community with 5000 iterations. Positive SES values indicate that the species group in question is more phylogenetically diverse than a random draw of the same number of species from the regional species pool (overdispersion). SES values above 1.96 will have P values above 0.975, indicating the SES’s statistical significance. Negative SES values indicate that the species group in question is less phylogenetically diverse than a random draw of the same number of species from the regional species pool (underdispersion). SES values below -1.96 will generate P values below 0.025, indicating the SES’s statistical significance.

### Weighting by abundance and range size

To test prediction 2, we calculated MPD and MNTD weighted by abundance and range size. We then compared the weighted versus unweighted standard effect sizes for each community. To weight these metrics by abundance, we used the “ses.mpd” function in ‘picante’ (Kembel et al., 2010), but switched the “abundance.weighted=” parameter from *false* to *true*. To weight by range size, we weighted by the log of the AOO value. Since it is directly correlated with species richness, PD cannot be weighted by abundance. We defined a significant change in diversity in response to weighting as a sign change in the SES or if SES absolute values became greater than 1.96.

### Removing species

To test prediction 3, we removed individual species and groups of species and recalculated phylogenetic diversity after the removal. For each site, we ranked species in order of decreasing rarity in terms of both local abundance and range size, such that the most abundant and largest range-size species were ranked first and the lowest abundance and smallest range-size species were ranked last. Then, we removed one species at a time in order of increasing rarity (removing the most abundant and largest range sizes first) and calculated the SES for PD, MPD, and MNTD of the resulting community. We also tested for autocorrelation across species of adjacent ranks within each site using the Durbin-Wallis test with 1, 2 and 3 steps to see if removal of similarly-ranked species had similar effects.

To test the impact of removing groups of species at a time, we removed groups of ten species in order of abundance and range size. Using a sliding window approach, we created groups of ten species, starting with the top 10 most abundant species, then shifting to species ranked 2-11, 3-12, and so on until the 10 least abundant species. We then calculated the SES of phylogenetic diversity for the community with each group of 10 removed. We analyzed all data in R version 4.3.1 (R Core Team, 2023).

## RESULTS

### Community composition & phylogenetic signal

The low elevation site contained 39 species across 17 families, the middle site had 27 species across 14 families, and the high site had 36 species across 20 families. We found that local abundance ranged from 0.5 to 573.5 individuals per site when pooled across plots and averaged across years, with a median average abundance of 10.5 individuals per site. At all three sites, the least abundant species had only one individual in one year with an average of 0.5 individuals per site across years (*Noccaea fendleri, Primula pauciflora* and *Rumex densiflorus* at the low elevation site*, Eucephalus breweri* at the middle and *Arctostaphylos uva-ursi, Erigeron speciosus, Hydrophyllum capitatum, Senecio integerrimus* and *Erigeron eatonii* at the high elevation site). The most abundant species at the low elevation site was *Collomia linearis* with an average abundance of 573.5. At the middle elevation site, *Heliomeris multiflora* was the most abundant with an average of 253.5 individuals, and at the high elevation site *Polygonum douglasii* was the most abundant with an average of 130 individuals. Range size measured as AOO spanned from 108 km^2^ for *Rumex densiflorus* to 469,436 km^2^ for *Achillea millefolium,* with a median range size 2316 km^2^. Both abundance and range size followed standard rank abundance curves (Fig. 1A-B).

**Figure. 1:**
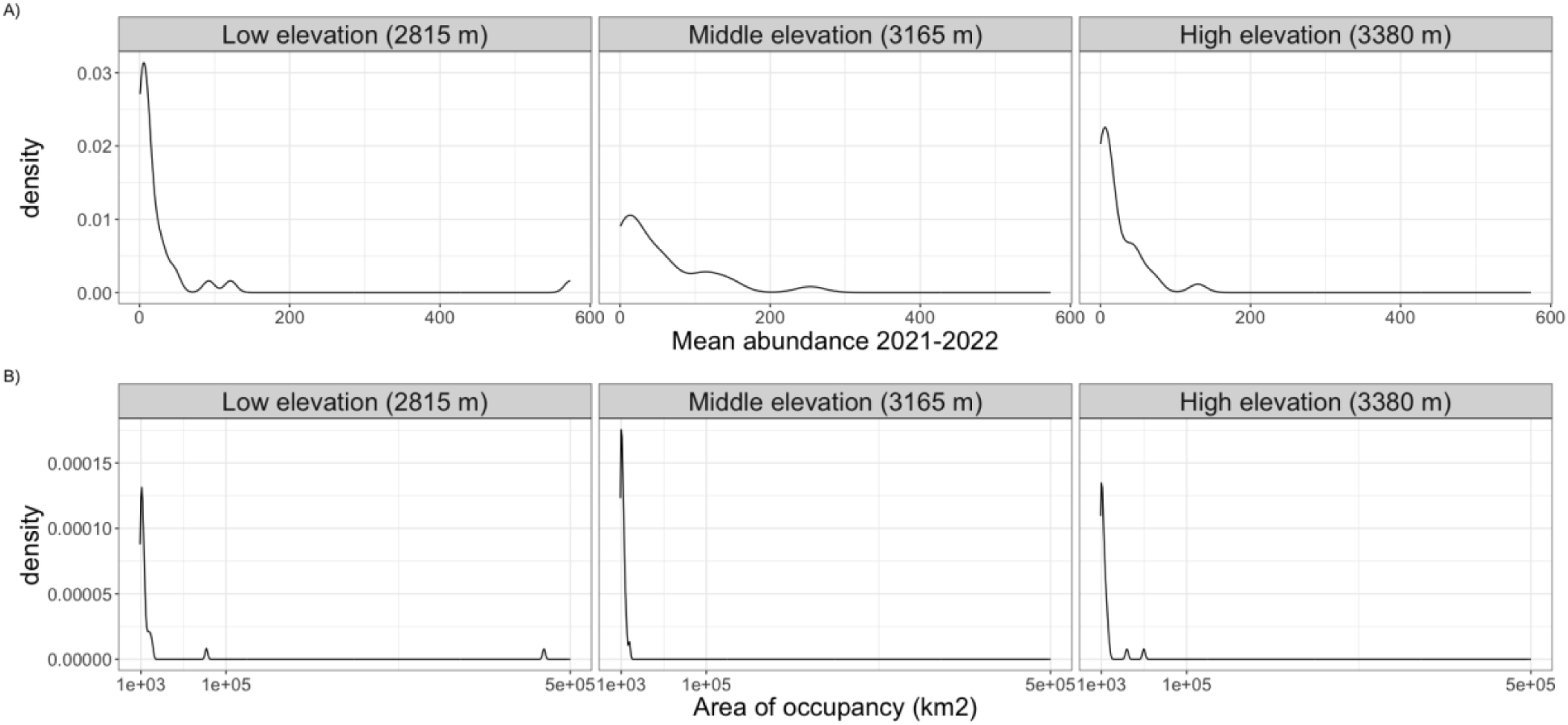
Rank abundance (A) and range size (B) curves for RMBL species at each site.

We found phylogenetic signal for abundance with only Blomberg’s K (Blomberg’s K = 0.304, P-value = 0.04; Pagel’s λ = 0.926, P-value =0.17; Fig. 2A), but not for range size with either metric (Blomberg’s K = 0.09, P-value = 0.7; Pagel’s λ = 7.3×10^−5^, P-value =1; Fig. 2B).

**Figure 2:**
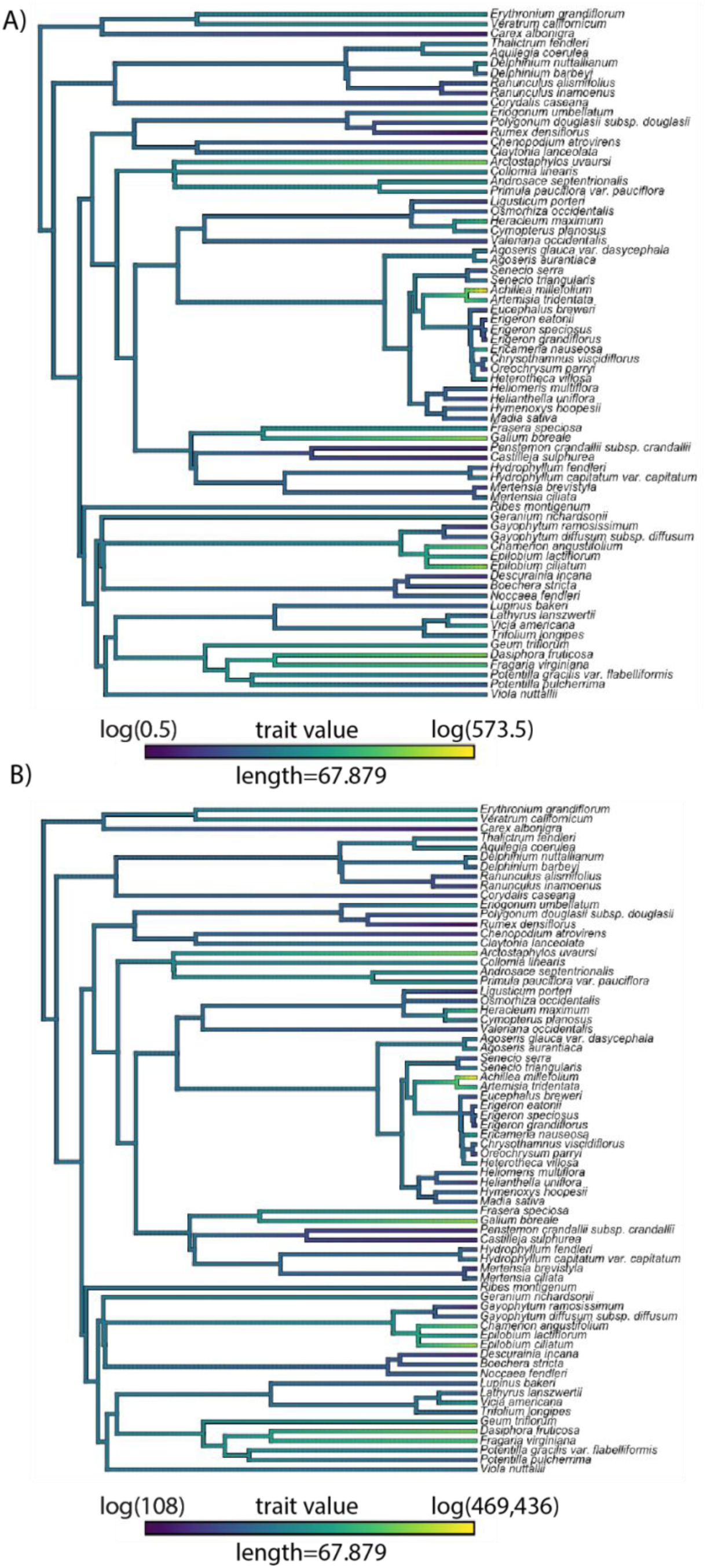
Phylogenies with log values of abundance (A) and range size (B) indicated by branch colors. A) Abundance shows significant phylogenetic signal for Blomberg’s K only (Blomberg’s K = 0.304, P-value = 0.04, Pagel’s λ = 0.926, P-value =0.168), while B) range size does not show any significant signal (Blomberg’s K = 0.09, P-value = 0.7, Pagel’s λ = 7.3×10^−5^, P-value =1).

### Weighting community phylogenetic metrics

Weighting MPD and MNTD by abundance (Fig. 3A) and range size (Fig. 3B) did not significantly change phylogenetic diversity at any of our sites. At the low elevation site, MPD SES was positive when unweighted, and negative when weighted by range size. However, both SES values were still very near zero and nonsignificant (Fig. 3B). Also at the low elevation site, weighting by abundance increased SES values for MPD and MNTD, but SES values stayed nonsignificant. At other sites, impacts of weighting were variable and depended on the metric in question (Fig. 3A-B), but never caused the SES values to be statistically significant.

**Figure 3:**
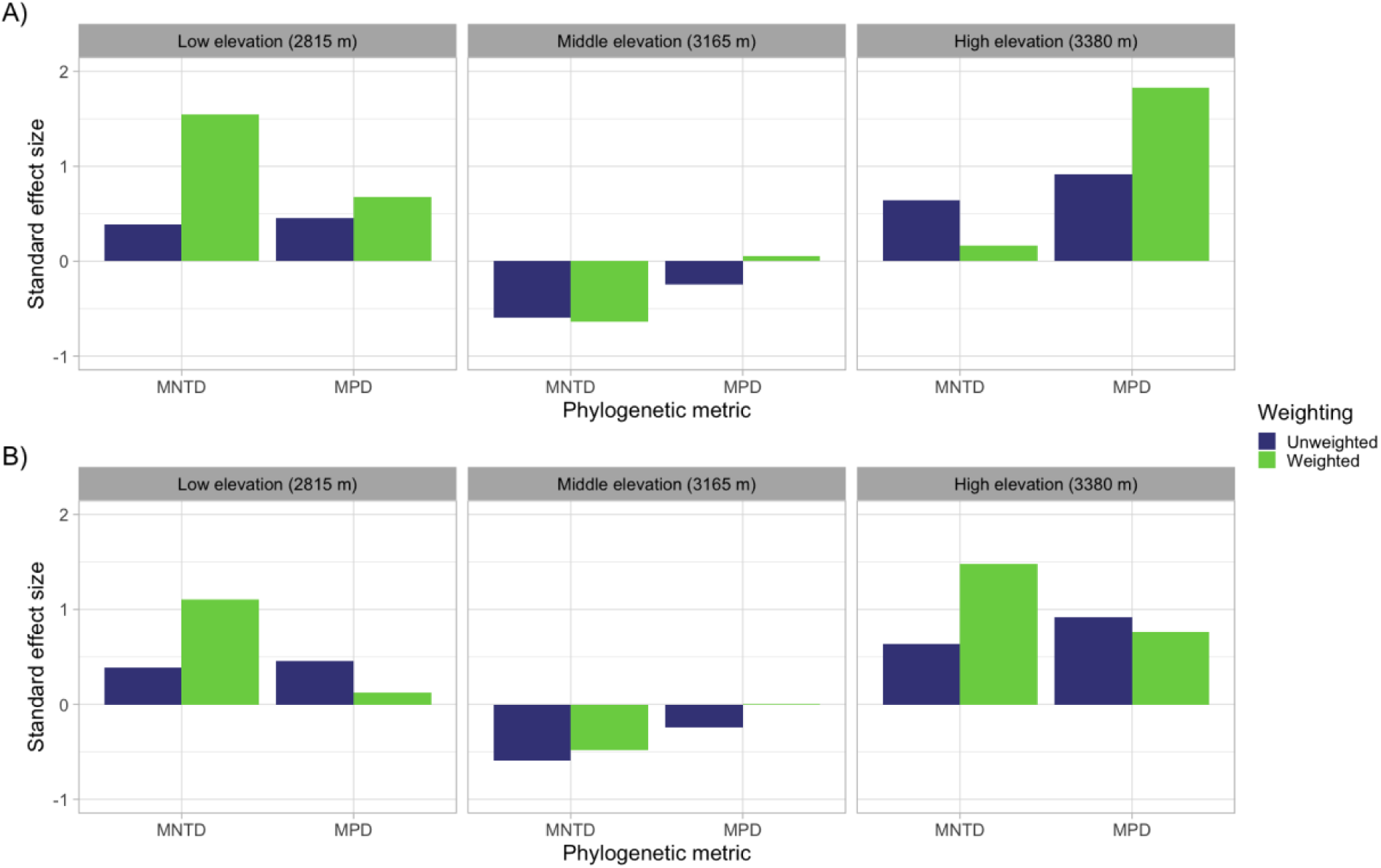
Standard effect sizes to compare weighted and unweighted MPD and MNTD. Unweighted values are in blue, and weighted values are in green. A) shows weighting by abundance. B) shows weighting by range size. Each of the three panels in A) and B) shows the different sites. Weighting by abundance and range size does not significantly change the sites’ MPD or MNTD.

### Removing species

Because range size did not show phylogenetic signal, we only show results for abundance (for single species range size removal results, see Appendix S3). For abundance, removing species in order of decreasing abundance impacted SES values inconsistently across metrics and sites (Fig. 4). Removing the rarest species did not consistently decrease phylogenetic diversity by any metric. At the high elevation site, removing the most abundant species did not change the SES for PD, slightly lowered SES for MPD, and increased the SES for MNTD (Fig. 4). Removing the least abundant species increased the SES for PD, MPD and MNTD, and the middle abundance species both increased and decreased SES values for all metrics depending on the species (Fig. 4).

**Figure 4:**
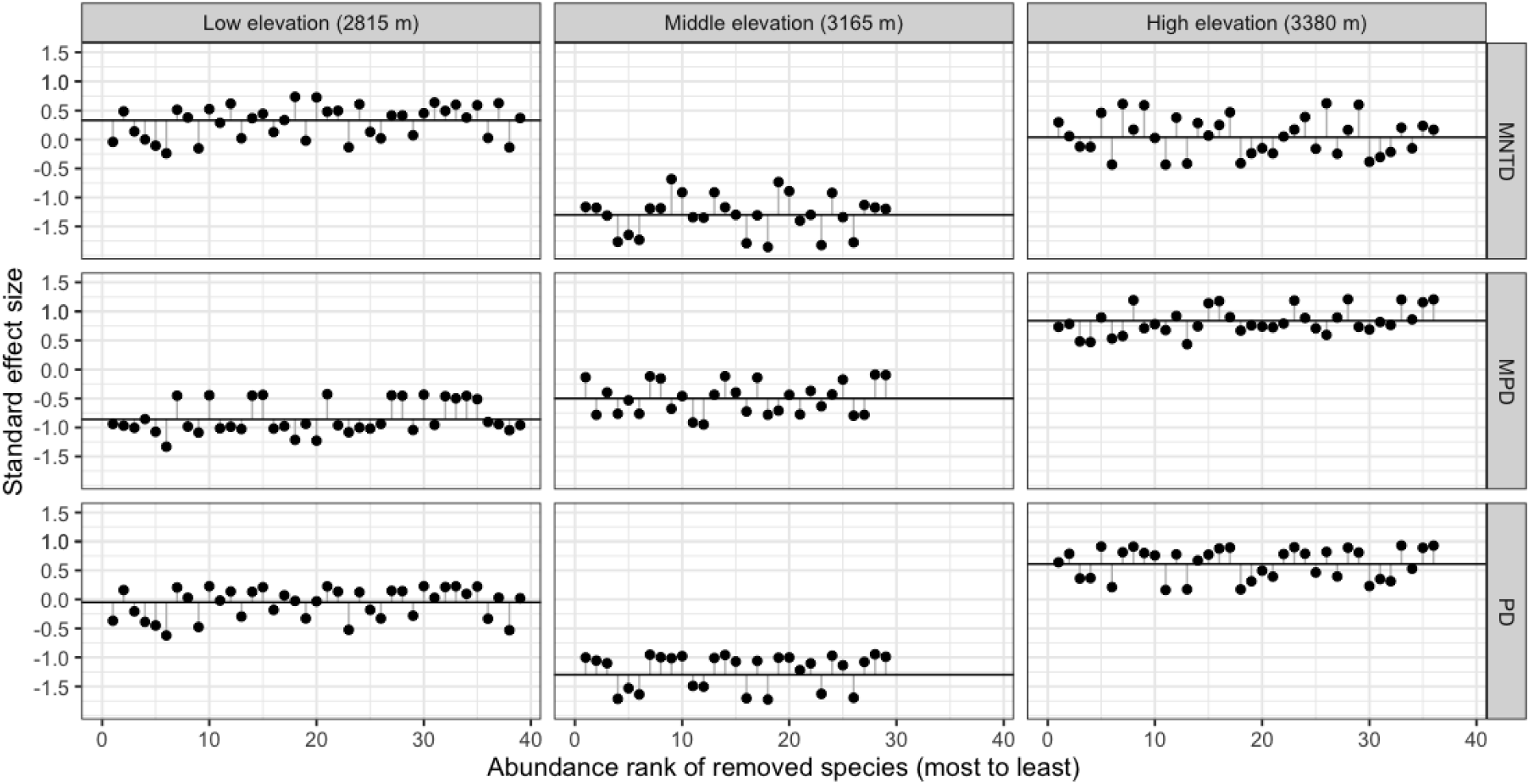
Change in SES values when individual species are removed in order of decreasing abundance. The horizontal black line in each figure shows the standard effect size of the metric for the overall community at each site, and the dots show how the SES changes when the species at the rank indicated on the x-axis is removed. MNTD values are shown in the top row, MPD values are in the middle row, and PD are on the bottom. The left column shows the low elevation site, the middle column is the middle, and the right is the high-elevation site. No clear pattern suggests that removing rarer species (right side of each panel) decreases MNTD, MPD or PD more than removing more abundant species (left side of each panel).

At the middle elevation site, removal of the most and least abundant species increased SES across all metrics (Fig. 4). Removing other low-abundance species (ranked 20-26 out of 29) both increased and decreased SES values depending on metric and individual species (Fig. 4). Like the high elevation site, removing the middle-ranked abundance species had mixed impacts on community phylogenetic diversity by all metrics (Fig. 4).

Finally, at the low elevation site removal of the most and least abundant species lowered SES values by all metrics (Fig. 4). For both PD and MTD, SES values decreased with the removal of each of the top five most abundant species (Fig. 4). This decrease in SES was also mostly true for MPD, except that the removal of the third most abundant species did not change the SES value. While the removal of the least abundant species decreased SES values for all metrics, removal of the next three least abundant species either did not change or increased SES values depending on metric and species (Fig. 4). Like the middle and high elevation sites, removing the middle ranked species at the low elevation site had variable impacts on SES values for all metrics (Fig. 4). We tested for autocorrelation between points within each site at one, two and three steps, and found no significant autocorrelation (P>0.05 for all; Appendix S4).

Removing groups of ten species in order of abundance also did not produce consistent patterns (Fig. 5). Removing the groups with the most rare species did not decrease SES for PD, and instead increased SES at all sites (Fig. 5). At the low elevation site, removing groups of common species had the most negative impact on SES values, while removing the most common species did not impact SES at the high elevation site, and increased the SES at the middle elevation site (Fig. 5). We only show results for PD here, as results for MPD and MNTD were similarly inconsistent (Appendix S5). Like the removal of individual species, we only removed groups of species based on abundance, because we found no phylogenetic signal in range size (Fig. 2B).

**Figure 5:**
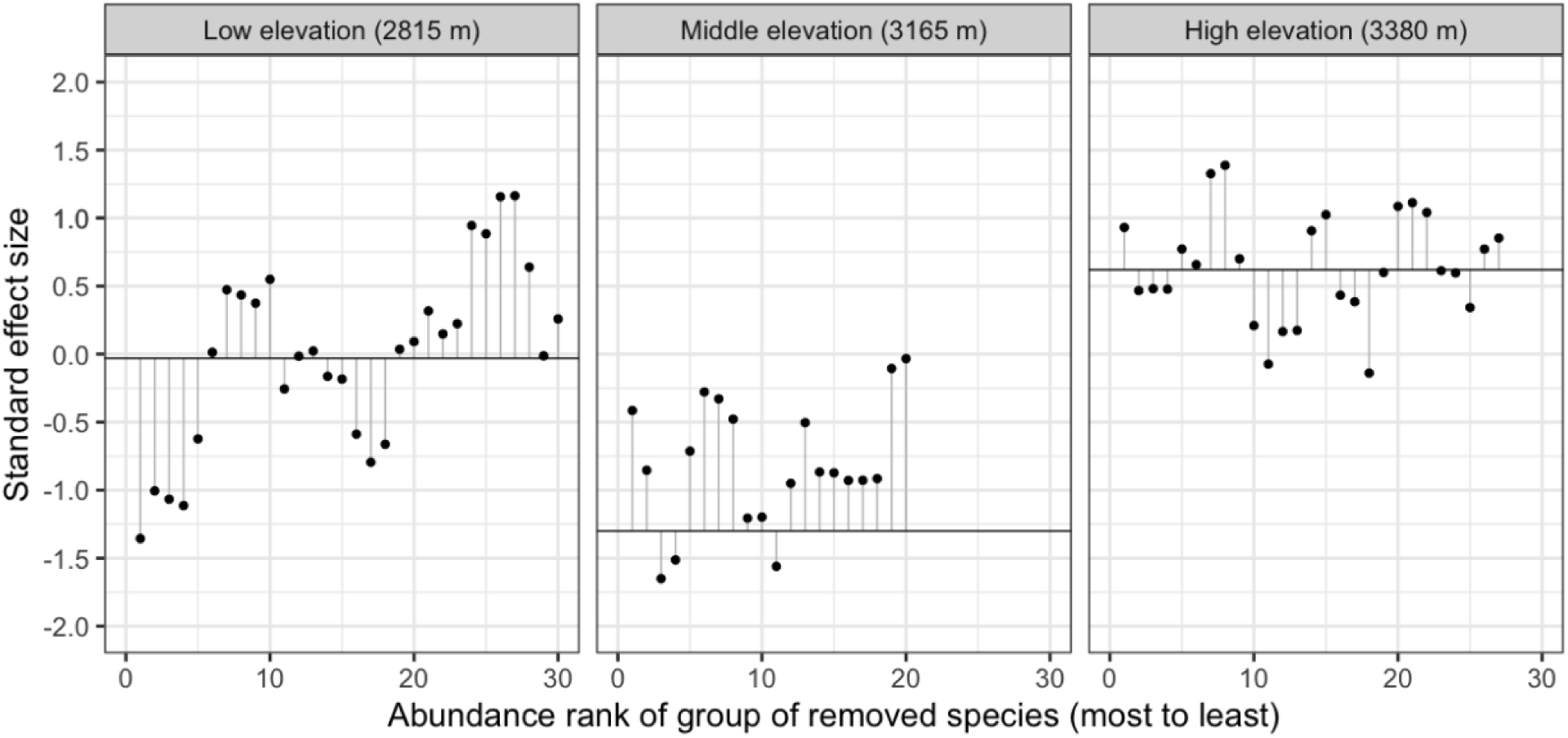
Change in SES values for PD when groups of ten species are removed in order of decreasing abundance. The horizontal black line in each figure shows the standard effect size of the metric for the overall community at each site, and dots show how the SES changes when the group is removed. Numbers on the x-axis rank the most abundant species in the group that was removed. For example, for the point at x=5, the species that were removed were ranked 5-14. The left panel shows the low-elevation site, middle panel is the middle elevation site, and right is the high elevation site. No clear pattern suggests that removing rarer species (right side of each panel) decreases PD more than removing more abundant species (left side of each panel).

## DISCUSSION

We hypothesized that rare species (quantified by local abundance and range size) would contribute disproportionately to phylogenetic diversity, but our results do not support our associated predictions. We found that abundance did show phylogenetic signal, but range size did not. We predicted that weighting MPD and MNTD by higher abundance and larger range size would decrease phylogenetic diversity (consistent with disproportionately high contributions by rare species), but it did not. Finally, we predicted that removing rare species (individually or in groups) would decrease phylogenetic diversity more than removing common species, but this was only sometimes the case. Our results are strikingly consistent in suggesting that rare species do not contribute disproportionately to phylogenetic diversity in this system.

Our results are consistent with a similar recent study that found that rare and common species contribute similarly to phylogenetic diversity in tallgrass prairies (Herzog and Latvis, 2022). The authors sampled 21 tallgrass prairie communities, and calculated PD each time they added increasingly rare species, which were quantified by percent cover. They then calculated breakpoints where phylogenetic diversity changed significantly from one species’ addition to the next. Still, they found that the location of statistically significant breakpoints was not consistently associated with rare species (Herzog and Latvis, 2022). While we did not calculate breakpoints in our data, Herzog and Latvis’s results are similar in that species’ phylogenetic contributions are not consistently predictable based on rarity.

Our results show no evidence that rare species are phylogenetically distant from the rest of the community. However, we did not investigate their functional uniqueness, which may produce a different pattern. Phylogenetic diversity often correlates poorly with variation in individual traits (CaraDonna and Inouye, 2015; Gerhold et al., 2015; Maitner et al., 2022; E-Vojtkó et al., 2023), indicating that phylogenetic and functional diversity patterns are not interchangeable. Similarly, prioritizing phylogenetic diversity for conservation efforts does not always maximize functional diversity, even when traits show phylogenetic signal (Mazel et al., 2017). Work in other systems has suggested that rare species provide the most unique functional trait combinations (Mouillot et al., 2013) and contribute significantly to functional diversity (Leitão et al., 2016; Coulon et al., 2023). Whether our rare RMBL plant community members are functionally unique or redundant is unknown, but our results suggest at least that low abundance and small range size do provide redundancy in terms of phylogenetic diversity.

In contrast to our results, Mi at al. (2012) tested for niche differentiation by comparing the phylogenetic diversity of rare and common species in temperate and tropical tree plots, and suggested that the niche differentiation hypothesis is supported in some of their plots because of rare species’ phylogenetic distinctiveness. While we now know we cannot assume relatedness predicts niche space in a straightforward way (Mason and Pavoine, 2013; Davies et al., 2016; Blonder, 2018), Mi and colleague’s results are informative. In particular, they found that rare species contribute more phylogenetic diversity only in plots with lower levels of disturbance (Mi et al., 2012). If disturbance has caused local extinctions of rare species before the time frame of our study, the rarest species will no longer be present, and we would instead be observing the less rare species that are still present. If this were the case, it may still be true that rare species historically contributed the most phylogenetic diversity, but we would not know. We also measured the impact of the removal of one species at a time and in groups of ten, but patterns of actual loss in this system are unknown and are likely to be highly system-specific (Zavaleta and Hulvey, 2004; Hooper et al., 2012; Sanders et al., 2018; Chase et al., 2020; Carmona et al., 2021). We did not evaluate disturbance or historical species loss at our sites, and this would be an important avenue for future research in this system.

The addition of new species via invasions or range shifts may also drive declines in phylogenetic diversity (Selvi et al., 2016; Fitt and Lancaster, 2017; Daru et al., 2021), regardless of whether native species are declining. We know that range shifting and invasive species are impacting this system at the same time others are declining (Renwick et al., 2016; Zorio et al., 2016; Shepard et al., 2022). We did not include non-native species in our analyses since quantifying their range size is more complex, and only four non-native species occur at low abundance in our plots to begin with. Three of the four non-native species are in Asteraceae, and as such would likely not change our results. Evidence does suggest plants are shifting their elevational ranges in this system (Zorio et al., 2016), however, so investigating the phylogenetic impact of newcomers may be important for future work.

We did find phylogenetic signal in abundance but not in range size. Herzog & Latvis (2022) tested for phylogenetic signals in abundance in their tall grass prairie communities and found signal at the regional scale and in 5 of their 21 communities. Previous work has tested phylogenetic signal in range size with mixed results (Morin and Lechowicz, 2013; Zacaï et al., 2017; Pigot et al., 2018). Pigot and colleagues (2018) found that introduced bird species have strong phylogenetic signal in their native range size, but random samples of birds do not show phylogenetic signal in range size, so this finding is unique to introduced species. Morin and Lechowicz (2013) found weak but significant signal in range size in tree species in forest plots worldwide. Zacaï and colleagues (2017) found that phylogenetic signal in range size varies regionally, and is likely influenced by spatio-temporal environmental stability over geological time. Each of these studies addresses range size across very broad spatial and temporal scales (Morin and Lechowicz, 2013; Zacaï et al., 2017; Pigot et al., 2018), while our study only includes present-day species native to the North American Rocky Mountains. This indicates that the scale tested likely influences whether or not range size shows phylogenetic signal, which is consistent with other scale-dependent community phylogenetic patterns (Cavender-Bares et al., 2009; Willis et al., 2009; Park et al., 2020).

Our findings suggest that rare species in this subalpine plant community do not disproportionately contribute to phylogenetic diversity, indicating that their evolutionary roles may be redundant with more common species. However, this raises important questions about their potential contributions to other dimensions of biodiversity, such as functional diversity and functional rarity. Violle et al. (2017) emphasize the critical role of functional rarity in maintaining ecosystem processes and resilience. Future research should explore whether the rare species in this system possess unique functional traits that contribute disproportionately to ecosystem functioning, even if they do not enhance phylogenetic diversity. Understanding the interplay between phylogenetic and functional rarity will help clarify the broader ecological significance of rare species and inform conservation strategies to preserve biodiversity and ecosystem stability in the face of global change. Integrating functional trait data with evolutionary frameworks would allow us to better assess whether rare species provide irreplaceable ecosystem functions or reinforce redundancy within ecological communities.

## CONCLUSIONS

Our study aimed to test whether rare species contribute disproportionately to phylogenetic diversity, a question central to understanding their ecological and evolutionary significance. Contrary to our hypothesis, we found that rare species in this subalpine plant community do not uniquely enhance phylogenetic diversity but instead may provide redundancy with more common species. These findings challenge assumptions that rare species universally enhance evolutionary diversity and raise questions about their broader ecological roles. We found that rare species do not contribute disproportionately to phylogenetic diversity and raise and lower phylogenetic diversity relative to richness depending on the identity of the species, elevation, and phylogenetic metric. This finding is consistent with previous work (Mi et al., 2012; Herzog and Latvis, 2022), and suggests that rare species may add phylogenetic redundancy rather than diversity in this community.

## ACKNOWLEDGEMENTS

This study was conducted on the traditional and unceded land of the Ute People. The authors thank RMBL and private landowners for land access, RMBL for securing permits, Dlugosch lab members for manuscript feedback, and Dan Park for discussing analyses and results. This work was funded by Rocky Mountain Biological Laboratory, Botanical Society of America, and University of Arizona Graduate & Professional Student Council awards to LNV and United States National Science Foundation grant #1750280 to KMD. BJE acknowledges that this material is based upon work supported by the U.S. Department of Energy, Office of Science, Office of Biological and Environmental Research, Earth and Environmental Systems Sciences Division and Data Management Program, under Award Number DEAC02-05CH11231, as part of the Watershed Function Scientific Focus Area and the ExaShed project. BJE was also supported by NSF awards 2225078 and 2225076.

## AUTHOR CONTRIBUTIONS

L.N.V. and K.M.D. conceived of the study. V.Z. collected field data. L.N.V. generated and analyzed range size data. L.N.V wrote the original draft with input from K.M.D. and V.Z. All authors provided feedback. L.N.V., B.J.E and K.M.D acquired funding. All authors approved the final version of the manuscript.

## DATA AVAILABILITY STATEMENT

Range size and abundance data can be accessed at https://doi.org/10.6073/pasta/320111b8e067c3b443fa25a719aac7ae. Code is available at https://github.com/lveldhuisen/phylogenetics-and-ranges.

Additional supporting information may be found online in the Supporting Information section at the end of the article.

**Appendix S1:** Citations for GBIF downloads used to calculate range size.

**Appendix S2:** Species in our research sites listed with replacement species used from Smith & Brown’s ALLMB.tre phylogeny (see also Methods).

**Appendix S3:** Results of phylogenetic diversity (PD, MPD & MNTD) from removing single species based on rage size.

**Appendix S4:** P-values for autocorrelation tests at each site for both abundance and range size, all three phylogenetic metrics and 1, 2 and 3 steps.

**Appendix S5:** Results of additional phylogenetic diversity metrics (MPD & MNTD) from removing groups of species based on abundance.

## LITERATURE CITED

Anderson, T. M., M. Lachance, and W. T. Starmer. 2004. The Relationship of Phylogeny to Community Structure: The Cactus Yeast Community. The American Naturalist 164: 709–721.

Blomberg, S. P., T. Garland, and A. R. Ives. 2003. Testing for Phylogenetic Signal in Comparative Data: Behavioral Traits Are More Labile. Evolution 57: 717–745.

Blonder, B. 2018. Hypervolume concepts in niche- and trait-based ecology. Ecography 41: 1441–1455.

Bracken, M. E. S., and N. H. N. Low. 2012. Realistic losses of rare species disproportionately impact higher trophic levels: Loss of rare ‘cornerstone’ species. Ecology letters 15: 461–467.

Cadotte, M. W. 2013. Experimental evidence that evolutionarily diverse assemblages result in higher productivity. Proceedings of the National Academy of Sciences of the United States of America 110: 8996–9000.

Cadotte, M. W., M. Carboni, X. Si, and S. Tatsumi. 2019. Do traits and phylogeny support congruent community diversity patterns and assembly inferences? The Journal of ecology 107: 2065–2077.

Cadotte, M. W., B. J. Cardinale, and T. H. Oakley. 2008. Evolutionary history and the effect of biodiversity on plant productivity. Proceedings of the National Academy of Sciences of the United States of America 105: 17012–17017.

Cadotte, M. W., J. Cavender-Bares, D. Tilman, and T. H. Oakley. 2009. Using phylogenetic, functional and trait diversity to understand patterns of plant community productivity. PloS one 4: e5695.

Cadotte, M. W., R. Dinnage, and D. Tilman. 2012. Phylogenetic diversity promotes ecosystem stability. Ecology 93: S223–S233.

CaraDonna, P. J., and D. W. Inouye. 2015. Phenological responses to climate change do not exhibit phylogenetic signal in a subalpine plant community. Ecology 96: 355–361.

Cardoso, P. 2017. red - an R package to facilitate species red list assessments according to the IUCN criteria. Biodiversity data journal: e20530.

Carmona, C. P., R. Tamme, M. Pärtel, F. de Bello, S. Brosse, P. Capdevila, R. González-M, et al. 2021. Erosion of global functional diversity across the tree of life. Science advances 7: eabf2675.

Cavender-Bares, J., K. H. Kozak, P. V. A. Fine, and S. W. Kembel. 2009. The merging of community ecology and phylogenetic biology. Ecology letters 12: 693–715.

Chamberlain, S. A., and C. Boettiger. 2017. R Python, and Ruby clients for GBIF species occurrence data. PeerJ Preprints.

Chase, J. M., S. A. Blowes, T. M. Knight, K. Gerstner, and F. May. 2020. Ecosystem decay exacerbates biodiversity loss with habitat loss. Nature 584: 238–243.

Chen, I.-C., J. K. Hill, R. Ohlemüller, D. B. Roy, and C. D. Thomas. 2011. Rapid range shifts of species associated with high levels of climate warming. Science 333: 1024–1026.

Coulon, N., M. Lindegren, E. Goberville, A. Toussaint, A. Receveur, and A. Auber. 2023. Threatened fish species in the Northeast Atlantic are functionally rare. Global ecology and biogeography: a journal of macroecology 32: 1827–1845.

Daru, B. H., T. J. Davies, C. G. Willis, E. K. Meineke, A. Ronk, M. Zobel, M. Pärtel, et al. 2021. Widespread homogenization of plant communities in the Anthropocene. Nature communications 12: 6983.

Davies, T. J., M. C. Urban, B. Rayfield, M. W. Cadotte, and P. R. Peres-Neto. 2016. Deconstructing the relationships between phylogenetic diversity and ecology: a case study on ecosystem functioning. Ecology 97: 2212–2222.

Dee, L. E., J. Cowles, F. Isbell, S. Pau, S. D. Gaines, and P. B. Reich. 2019. When Do Ecosystem Services Depend on Rare Species? Trends in ecology & evolution 34: 746–758.

Delavaux, C. S., T. W. Crowther, C. M. Zohner, N. M. Robmann, T. Lauber, J. van den Hoogen, S. Kuebbing, et al. 2023. Native diversity buffers against severity of non-native tree invasions. Nature 621: 773–781.

Dinnage, R., M. W. Cadotte, N. M. Haddad, G. M. Crutsinger, and D. Tilman. 2012. Diversity of plant evolutionary lineages promotes arthropod diversity. Ecology letters 15: 1308–1317.

Eiserhardt, W. L., F. Borchsenius, C. M. Plum, A. Ordonez, and J.-C. Svenning. 2015. Climate-driven extinctions shape the phylogenetic structure of temperate tree floras. Ecology letters 18: 263–272.

Enquist, B. J., X. Feng, B. Boyle, B. Maitner, E. A. Newman, P. M. Jørgensen, P. R. Roehrdanz, et al. 2019. The commonness of rarity: Global and future distribution of rarity across land plants. Science Advances 5: eaaz0414.

E-Vojtkó, A., F. de Bello, Z. Lososová, and L. Götzenberger. 2023. Phylogenetic diversity is a weak proxy for functional diversity but they are complementary in explaining community assembly patterns in temperate vegetation. The Journal of ecology.

Faith, D. P. 1992. Conservation evaluation and phylogenetic diversity. Biological conservation 61: 1–10.

Fitt, R. N. L., and L. T. Lancaster. 2017. Range shifting species reduce phylogenetic diversity in high latitude communities via competition. The Journal of animal ecology 86: 543–555.

Galland, T., G. Adeux, H. Dvořáková, A. E-Vojtkó, I. Orbán, M. Lussu, J. Puy, et al. 2019. Colonization resistance and establishment success along gradients of functional and phylogenetic diversity in experimental plant communities. The Journal of ecology 107: 2090–2104.

Gaston, K. J. 1994. Biodiversity - measurement. Progress in Physical Geography: Earth and Environment 18: 565–574.

Gaston, K. J., and R. A. Fuller. 2009. The sizes of species’ geographic ranges. The Journal of applied ecology 46: 1–9.

Gerhold, P., J. F. Cahill, M. Winter, I. V. Bartish, and A. Prinzing. 2015. Phylogenetic patterns are not proxies of community assembly mechanisms (they are far better). Functional ecology 29: 600–614.

Ge, X.-Y. M., J. P. Scholl, U. Basinger, T. E. Huxman, and D. L. Venable. 2019. Functional trait trade-off and species abundance: insights from a multi-decadal study. Ecology letters 22: 583–592.

González-Orozco, C. E., L. J. Pollock, A. H. Thornhill, B. D. Mishler, N. Knerr, S. W. Laffan, J. T. Miller, et al. 2016. Phylogenetic approaches reveal biodiversity threats under climate change. Nature climate change 6: 1110–1114.

Grime, J. P. 1998. Benefits of plant diversity to ecosystems: immediate, filter and founder effects. The Journal of ecology 86: 902–910.

Hanski, I. 1982. Dynamics of Regional Distribution: The Core and Satellite Species Hypothesis. Oikos 38: 210–221.

Hardie, D. C., and J. A. Hutchings. 2010. Evolutionary ecology at the extremes of species’ ranges. Environmental Review 18: 1–20.

Harnik, P. G., C. Simpson, and J. L. Payne. 2012. Long-term differences in extinction risk among the seven forms of rarity. Proceedings. Biological sciences / The Royal Society 279: 4969–4976.

Herzog, S. A., and M. Latvis. 2022. Community-level phylogenetic diversity does not differ between rare and common lineages across tallgrass prairies in the northern Great Plains. Ecology and evolution 12: e9453.

Hooper, D. U., E. C. Adair, B. J. Cardinale, J. E. K. Byrnes, B. A. Hungate, K. L. Matulich, A. Gonzalez, et al. 2012. A global synthesis reveals biodiversity loss as a major driver of ecosystem change. Nature 486: 105–108.

Huang, M., X. Liu, M. W. Cadotte, and S. Zhou. 2020. Functional and phylogenetic diversity explain different components of diversity effects on biomass production. Oikos 129: 1185–1195.

Jiao, S., W. Chen, and G. Wei. 2017. Biogeography and ecological diversity patterns of rare and abundant bacteria in oil-contaminated soils. Molecular ecology 26: 5305–5317.

Kembel, S. W., P. D. Cowan, M. R. Helmus, W. K. Cornwell, H. Morlon, D. D. Ackerly, S. P. Blomberg, and C. O. Webb. 2010. Picante: R tools for integrating phylogenies and ecology. Bioinformatics 26: 1463–1464.

Leitão, R. P., J. Zuanon, S. Villéger, S. E. Williams, C. Baraloto, C. Fortunel, F. P. Mendonça, and D. Mouillot. 2016. Rare species contribute disproportionately to the functional structure of species assemblages. Proceedings. Biological sciences / The Royal Society 283.

Maitner, B. S., D. S. Park, B. J. Enquist, and K. M. Dlugosch. 2022. Where we’ve been and where we’re going: the importance of source communities in predicting establishment success from phylogenetic relationships. Ecography 2022.

Manes, S., M. J. Costello, H. Beckett, A. Debnath, E. Devenish-Nelson, K.-A. Grey, R. Jenkins, et al. 2021. Endemism increases species’ climate change risk in areas of global biodiversity importance. Biological conservation 257: 109070.

Mason, N. W. H., and S. Pavoine. 2013. Does trait conservatism guarantee that indicators of phylogenetic community structure will reveal niche-based assembly processes along stress gradients? Journal of vegetation science: official organ of the International Association for Vegetation Science 24: 820–833.

Mazel, F., A. O. Mooers, G. V. D. Riva, and M. W. Pennell. 2017. Conserving Phylogenetic Diversity Can Be a Poor Strategy for Conserving Functional Diversity. Systematic biology 66: 1019–1027.

Milcu, A., E. Allan, C. Roscher, T. Jenkins, S. T. Meyer, D. Flynn, H. Bessler, et al. 2013. Functionally and phylogenetically diverse plant communities key to soil biota. Ecology 94: 1878–1885.

Mi, X., N. G. Swenson, R. Valencia, W. J. Kress, D. L. Erickson, Á. J. Pérez, H. Ren, et al. 2012. The contribution of rare species to community phylogenetic diversity across a global network of forest plots. The American naturalist 180: E17–30.

Molina-Venegas, R., M. Á. Rodríguez, M. Pardo-de-Santayana, C. Ronquillo, and D. J. Mabberley. 2021. Maximum levels of global phylogenetic diversity efficiently capture plant services for humankind. Nature ecology & evolution 5: 583–588.

Morin, X., and M. J. Lechowicz. 2013. Niche breadth and range area in North American trees. Ecography 36: 300–312.

Mouillot, D., D. R. Bellwood, C. Baraloto, J. Chave, R. Galzin, M. Harmelin-Vivien, M. Kulbicki, et al. 2013. Rare species support vulnerable functions in high-diversity ecosystems. PLoS biology 11: e1001569.

Pagel, M. 1999. Inferring the historical patterns of biological evolution. Nature 401: 877–884.

Park, D. S., X. Feng, B. S. Maitner, K. C. Ernst, and B. J. Enquist. 2020. Darwin’s naturalization conundrum can be explained by spatial scale. Proceedings of the National Academy of Sciences of the United States of America 117: 10904–10910.

Pelser, P. B., B. Nordenstam, J. W. Kadereit, and L. E. Watson. 2007. An ITS phylogeny of tribe Senecioneae (Asteraceae) and a new delimitation of Senecio L. Taxon 56: 1077–1104.

Pigot, A. L., E. E. Dyer, D. W. Redding, P. Cassey, G. H. Thomas, and T. M. Blackburn. 2018. Species invasions and the phylogenetic signal in geographical range size. Global ecology and biogeography: a journal of macroecology 27: 1080–1092.

Rabinowitz, D. 1981. Seven forms of rarity. Biological aspects of rare plant conservation.

R Core Team. 2023. R: A language and environment for statistical computing.

Renwick, K. M., M. E. Rocca, and T. J. Stohlgren. 2016. Biotic disturbance facilitates range shift at the trailing but not the leading edge of lodgepole pine’s altitudinal distribution. Journal of vegetation science: official organ of the International Association for Vegetation Science 27: 780–788.

Revell, L. J. 2011. phytools: an R package for phylogenetic comparative biology (and other things). Methods in Ecology & Evolution 3: 217–223.

Sanders, D., E. Thébault, R. Kehoe, and F. J. Frank van Veen. 2018. Trophic redundancy reduces vulnerability to extinction cascades. Proceedings of the National Academy of Sciences of the United States of America 115: 2419–2424.

Selvi, F., E. Carrari, and A. Coppi. 2016. Impact of pine invasion on the taxonomic and phylogenetic diversity of a relict Mediterranean forest ecosystem. Forest ecology and management 367: 1–11.

Sexton, J. P., P. J. McIntyre, A. L. Angert, and K. J. Rice. 2009. Evolution and Ecology of Species Range Limits. Annual review of ecology, evolution, and systematics 40: 415–436.

Shepard, I. D., S. A. Wissinger, Z. T. Wood, and H. S. Greig. 2022. Predators balance consequences of climate-change-induced habitat shifts for range-shifting and resident species. The Journal of animal ecology 91: 334–344.

Sheth, S. N., and A. L. Angert. 2018. Demographic compensation does not rescue populations at a trailing range edge. Proceedings of the National Academy of Sciences of the United States of America 115: 2413–2418.

Sloat, L. L., A. N. Henderson, C. Lamanna, and B. J. Enquist. 2015. The Effect of the Foresummer Drought on Carbon Exchange in Subalpine Meadows. Ecosystems 18: 533–545.

Smith, S. A., and J. W. Brown. 2018. Constructing a broadly inclusive seed plant phylogeny. American journal of botany 105: 302–314.

Staude, I. R., D. M. Waller, M. Bernhardt-Römermann, A. D. Bjorkman, J. Brunet, P. De Frenne, R. Hédl, et al. 2020. Replacements of small-by large-ranged species scale up to diversity loss in Europe’s temperate forest biome. Nature ecology & evolution 4: 802–808.

Tucker, C. M., M. W. Cadotte, S. B. Carvalho, T. J. Davies, S. Ferrier, S. A. Fritz, R. Grenyer, et al. 2017. A guide to phylogenetic metrics for conservation, community ecology and macroecology. Biological reviews of the Cambridge Philosophical Society 92: 698–715.

Tucker, C. M., M. W. Cadotte, T. J. Davies, and T. G. Rebelo. 2012. Incorporating geographical and evolutionary rarity into conservation prioritization. Conservation biology: the journal of the Society for Conservation Biology 26: 593–601.

Veldhuisen, L. N., B. J. Enquist, and K. M. Dlugosch. 2023. Phylogenetic diversity of flowering plants declines throughout the growing season across a subalpine elevational gradient. bioRxiv: 2023.11.06.565878.

Violle, C., W. Thuiller, N. Mouquet, F. Munoz, N. J. B. Kraft, M. W. Cadotte, S. W. Livingstone, and D. Mouillot. 2017. Functional rarity: The ecology of outliers. Trends in ecology & evolution 32: 356–367.

White, H. J., C. M. McKeon, R. J. Pakeman, and Y. M. Buckley. 2023. The contribution of geographically common and rare species to the spatial distribution of biodiversity. Global ecology and biogeography: a journal of macroecology 32: 1730–1747.

Willis, C. G., M. Halina, C. Lehman, P. B. Reich, A. Keen, S. McCarthy, and J. Cavender-Bares. 2009. Phylogenetic community structure in Minnesota oak savanna is influenced by spatial extent and environmental variation. Ecography 33: no–no.

Willis, C. G., B. Ruhfel, R. B. Primack, A. J. Miller-Rushing, and C. C. Davis. 2008. Phylogenetic patterns of species loss in Thoreau’s woods are driven by climate change. Proceedings of the National Academy of Sciences 105: 17029–17033.

Yessoufou, K., and T. J. Davies. 2016. Reconsidering the Loss of Evolutionary History: How Does Non-random Extinction Prune the Tree-of-Life? *In* R. Pellens, and P. Grandcolas [eds.], Biodiversity Conservation and Phylogenetic Systematics: Preserving our evolutionary heritage in an extinction crisis, Topics in Biodiversity and Conservation, 57–80. Springer International Publishing, Cham.

Zacaï, A., E. Fara, A. Brayard, R. Laffont, J.-L. Dommergues, and C. Meister. 2017. Phylogenetic conservatism of species range size is the combined outcome of phylogeny and environmental stability. Journal of biogeography 44: 2451–2462.

Zavaleta, E. S., and K. B. Hulvey. 2004. Realistic species losses disproportionately reduce grassland resistance to biological invaders. *Science (New York*, N.Y.) 306: 1175–1177.

Zorio, S. D., C. F. Williams, and K. A. Aho. 2016. Sixty-Five Years of Change in Montane Plant Communities in Western Colorado, U.S.A. Arctic, Antarctic, and Alpine Research 48: 703–722.

