## Appendix S1 for "Rare species do not disproportionately contribute to phylogenetic diversity in a subalpine plant community"

Table S1: Citations for all GBIF downloads used to calculate range size.

| Species | GBIF citation |
|---|---|
| *Achillea millefolium* | GBIF Occurrence Download https://doi.org/10.15468/dl.ngwep5 Accessed from R via rgbif (https://github.com/ropensci/rgbif) on 2023-11-02 |
| *Agoseris aurantiaca* | GBIF Occurrence Download https://doi.org/10.15468/dl.f8st3x Accessed from R via rgbif (https://github.com/ropensci/rgbif) on 2023-10-12 |
| *Agoseris glauca* | GBIF Occurrence Download https://doi.org/10.15468/dl.uaaa54 Accessed from R via rgbif (https://github.com/ropensci/rgbif) on 2023-10-17 |
| *Androsace septentrionalis* | GBIF Occurrence Download https://doi.org/10.15468/dl.7vb3c9 Accessed from R via rgbif (https://github.com/ropensci/rgbif) on 2023-10-17 |
| *Aquilegia coerulea* | GBIF Occurrence Download https://doi.org/10.15468/dl.kxwjam Accessed from R via rgbif (https://github.com/ropensci/rgbif) on 2023-10-24 |
| *Arctostaphylos uva-ursi* | GBIF Occurrence Download https://doi.org/10.15468/dl.8app72 Accessed from R via rgbif (https://github.com/ropensci/rgbif) on 2024-10-28 |
| *Artemisia tridentata* | GBIF Occurrence Download https://doi.org/10.15468/dl.zj2pyd Accessed from R via rgbif (https://github.com/ropensci/rgbif) on 2023-10-24 |
| *Boechera stricta* | GBIF Occurrence Download https://doi.org/10.15468/dl.e6npex Accessed from R via rgbif (https://github.com/ropensci/rgbif) on 2023-10-27 |
| *Carex albonigra* | GBIF Occurrence Download https://doi.org/10.15468/dl.8u7b56 Accessed from R via rgbif (https://github.com/ropensci/rgbif) on 2023-10-27 |
| *Castilleja sulphurea* | GBIF Occurrence Download https://doi.org/10.15468/dl.gkcabc Accessed from R via rgbif (https://github.com/ropensci/rgbif) on 2023-10-27 |
| *Chamerion angustifolium* | GBIF Occurrence Download https://doi.org/10.15468/dl.c4px5r Accessed from R via rgbif (https://github.com/ropensci/rgbif) on 2023-10-27 |
| *Chenopodium atrovirens* | GBIF Occurrence Download https://doi.org/10.15468/dl.d8ntph Accessed from R via rgbif (https://github.com/ropensci/rgbif) on 2023-10-31 |
| *Chrysothamnus viscidiflorus* | GBIF Occurrence Download https://doi.org/10.15468/dl.dtqvzy Accessed from R via rgbif (https://github.com/ropensci/rgbif) on 2023-10-27 |
| *Claytonia lanceolata* | GBIF Occurrence Download https://doi.org/10.15468/dl.tjqnhn Accessed from R via rgbif (https://github.com/ropensci/rgbif) on 2023-10-27 |
| *Collomia linearis* | GBIF Occurrence Download https://doi.org/10.15468/dl.emwjeg Accessed from R via rgbif (https://github.com/ropensci/rgbif) on 2023-10-26 |
| *Corydalis caseana* | GBIF Occurrence Download https://doi.org/10.15468/dl.fy22a9 Accessed from R via rgbif (https://github.com/ropensci/rgbif) on 2023-10-31 |

| | |
|---|---|
| *Cymopterus lemmonii* | GBIF Occurrence Download https://doi.org/10.15468/dl.rb9feq Accessed from R via rgbif (https://github.com/ropensci/rgbif) on 2023-10-31 |
| *Dasiphora fruticosa* | GBIF Occurrence Download https://doi.org/10.15468/dl.9s64m4 Accessed from R via rgbif (https://github.com/ropensci/rgbif) on 2023-10-31 |
| *Delphinium barbeyi* | GBIF Occurrence Download https://doi.org/10.15468/dl.dqhdwy Accessed from R via rgbif (https://github.com/ropensci/rgbif) on 2023-10-31 |
| *Delphinium nuttallianum* | GBIF Occurrence Download https://doi.org/10.15468/dl.c2t5hq Accessed from R via rgbif (https://github.com/ropensci/rgbif) on 2023-10-31 |
| *Descurainia sp.* | GBIF Occurrence Download https://doi.org/10.15468/dl.2hfuyy Accessed from R via rgbif (https://github.com/ropensci/rgbif) on 2023-10-31 |
| *Epilobium ciliatum* | GBIF Occurrence Download https://doi.org/10.15468/dl.ysw44u Accessed from R via rgbif (https://github.com/ropensci/rgbif) on 2024-10-28 |
| *Epilobium lactiflorum* | GBIF Occurrence Download https://doi.org/10.15468/dl.2knuvb Accessed from R via rgbif (https://github.com/ropensci/rgbif) on 2024-10-28 |
| *Ericameria nauseosa* | GBIF Occurrence Download https://doi.org/10.15468/dl.zdybek Accessed from R via rgbif (https://github.com/ropensci/rgbif) on 2023-10-31 |
| *Erigeron eatonii* | GBIF Occurrence Download https://doi.org/10.15468/dl.pc4pjw Accessed from R via rgbif (https://github.com/ropensci/rgbif) on 2024-10-29 |
| *Erigeron elatior* | GBIF Occurrence Download https://doi.org/10.15468/dl.qgkq9b Accessed from R via rgbif (https://github.com/ropensci/rgbif) on 2024-10-29 |
| *Erigeron speciosus* | GBIF Occurrence Download https://doi.org/10.15468/dl.2h3vkh Accessed from R via rgbif (https://github.com/ropensci/rgbif) on 2023-10-31 |
| *Eriogonum umbellatum* | GBIF Occurrence Download https://doi.org/10.15468/dl.kzyspv Accessed from R via rgbif (https://github.com/ropensci/rgbif) on 2023-10-30 |
| *Erythronium grandiflorum* | GBIF Occurrence Download https://doi.org/10.15468/dl.3pxvek Accessed from R via rgbif (https://github.com/ropensci/rgbif) on 2023-10-30 |
| *Eucephalus engelmannii* | GBIF Occurrence Download https://doi.org/10.15468/dl.e5t4cd Accessed from R via rgbif (https://github.com/ropensci/rgbif) on 2023-10-31 |
| *Fragaria virginiana* | GBIF Occurrence Download https://doi.org/10.15468/dl.md3pnw Accessed from R via rgbif (https://github.com/ropensci/rgbif) on 2023-11-01 |
| *Frasera speciosa* | GBIF Occurrence Download https://doi.org/10.15468/dl.ptvhem Accessed from R via rgbif (https://github.com/ropensci/rgbif) on 2023-11-01 |
| *Galium boreale* | GBIF Occurrence Download https://doi.org/10.15468/dl.3nbsmy Accessed from R via rgbif (https://github.com/ropensci/rgbif) on 2024-10-28 |
| *Gayophytum diffusum* | GBIF Occurrence Download https://doi.org/10.15468/dl.emajxk Accessed from R via rgbif (https://github.com/ropensci/rgbif) on 2023-11-01 |
| *Gayophytum ramosissimum* | GBIF Occurrence Download https://doi.org/10.15468/dl.h4ueaq Accessed from R via rgbif (https://github.com/ropensci/rgbif) on 2024-10-28 |

| | |
|---|---|
| *Geranium richardsonii* | GBIF Occurrence Download https://doi.org/10.15468/dl.3sc2qk Accessed from R via rgbif (https://github.com/ropensci/rgbif) on 2023-11-01 |
| *Geum triflorum* | GBIF Occurrence Download https://doi.org/10.15468/dl.qbt264 Accessed from R via rgbif (https://github.com/ropensci/rgbif) on 2023-11-02 |
| *Helianthella quinquenervis* | GBIF Occurrence Download https://doi.org/10.15468/dl.2zsm99 Accessed from R via rgbif (https://github.com/ropensci/rgbif) on 2023-11-02 |
| *Heliomeris multiflora* | GBIF Occurrence Download https://doi.org/10.15468/dl.7w7ekx Accessed from R via rgbif (https://github.com/ropensci/rgbif) on 2023-11-02 |
| *Heracleum maximum* | GBIF Occurrence Download https://doi.org/10.15468/dl.gudr5j Accessed from R via rgbif (https://github.com/ropensci/rgbif) on 2024-10-28 |
| *Heterotheca villosa* | GBIF Occurrence Download https://doi.org/10.15468/dl.mwvzf7 Accessed from R via rgbif (https://github.com/ropensci/rgbif) on 2023-11-02 |
| *Hydrophyllum capitatum* | GBIF Occurrence Download https://doi.org/10.15468/dl.243cdf Accessed from R via rgbif (https://github.com/ropensci/rgbif) on 2023-11-01 |
| *Hydrophyllum fendleri* | GBIF Occurrence Download https://doi.org/10.15468/dl.wjy5s4 Accessed from R via rgbif (https://github.com/ropensci/rgbif) on 2023-11-02 |
| *Hymenoxys hoopesii* | GBIF Occurrence Download https://doi.org/10.15468/dl.bnht3v Accessed from R via rgbif (https://github.com/ropensci/rgbif) on 2023-11-08 |
| *Lathyrus lanszwertii* | GBIF Occurrence Download https://doi.org/10.15468/dl.btabfm Accessed from R via rgbif (https://github.com/ropensci/rgbif) on 2023-11-02 |
| *Ligusticum porteri* | GBIF Occurrence Download https://doi.org/10.15468/dl.s94489 Accessed from R via rgbif (https://github.com/ropensci/rgbif) on 2023-11-01 |
| *Lupinus bakeri* | GBIF Occurrence Download https://doi.org/10.15468/dl.ja8skt Accessed from R via rgbif (https://github.com/ropensci/rgbif) on 2023-11-03 |
| *Madia glomerata* | GBIF Occurrence Download https://doi.org/10.15468/dl.fqy5zj Accessed from R via rgbif (https://github.com/ropensci/rgbif) on 2023-11-03 |
| *Mertensia brevistyla* | GBIF Occurrence Download https://doi.org/10.15468/dl.u63zw9 Accessed from R via rgbif (https://github.com/ropensci/rgbif) on 2023-11-01 |
| *Mertensia ciliata* | GBIF Occurrence Download https://doi.org/10.15468/dl.r2gvzp Accessed from R via rgbif (https://github.com/ropensci/rgbif) on 2023-11-04 |
| *Noccaea fendleri* | GBIF Occurrence Download https://doi.org/10.15468/dl.qp84jx Accessed from R via rgbif (https://github.com/ropensci/rgbif) on 2023-11-0 |
| *Oreochrysum parryi* | GBIF Occurrence Download https://doi.org/10.15468/dl.6emwp9 Accessed from R via rgbif (https://github.com/ropensci/rgbif) on 2023-11-03 |
| *Osmorhiza occidentalis* | GBIF Occurrence Download https://doi.org/10.15468/dl.ks82fm Accessed from R via rgbif (https://github.com/ropensci/rgbif) on 2023-11-03 |
| *Penstemon crandallii* | GBIF Occurrence Download https://doi.org/10.15468/dl.aqyyk9 Accessed from R via rgbif (https://github.com/ropensci/rgbif) on 2024-10-28 |

| | |
|---|---|
| *Polygonum douglasii* | GBIF Occurrence Download https://doi.org/10.15468/dl.a2d2by Accessed from R via rgbif (https://github.com/ropensci/rgbif) on 2023-11-04 |
| *Potentilla gracilis* | GBIF Occurrence Download https://doi.org/10.15468/dl.8nfm36 Accessed from R via rgbif (https://github.com/ropensci/rgbif) on 2023-11-05 |
| *Potentilla pulcherrima* | GBIF Occurrence Download https://doi.org/10.15468/dl.zp66u2 Accessed from R via rgbif (https://github.com/ropensci/rgbif) on 2024-10-29 |
| *Primula pauciflora* | GBIF Occurrence Download https://doi.org/10.15468/dl.7fvb56 Accessed from R via rgbif (https://github.com/ropensci/rgbif) on 2023-11-03 |
| *Ranunculus alismifolius* | GBIF Occurrence Download https://doi.org/10.15468/dl.m8bjhy Accessed from R via rgbif (https://github.com/ropensci/rgbif) on 2023-11-03 |
| *Ranunculus inamoenus* | GBIF Occurrence Download https://doi.org/10.15468/dl.fanvp2 Accessed from R via rgbif (https://github.com/ropensci/rgbif) on 2023-11-03 |
| *Ribes montigenum* | GBIF Occurrence Download https://doi.org/10.15468/dl.wrvtqm Accessed from R via rgbif (https://github.com/ropensci/rgbif) on 2023-11-03 |
| *Rumex densiflorus* | GBIF Occurrence Download https://doi.org/10.15468/dl.ze396f Accessed from R via rgbif (https://github.com/ropensci/rgbif) on 2024-10-28 |
| *Senecio serra* | GBIF Occurrence Download https://doi.org/10.15468/dl.g8kz8b Accessed from R via rgbif (https://github.com/ropensci/rgbif) on 2023-11-04 |
| *Senecio integerrimus* | GBIF Occurrence Download https://doi.org/10.15468/dl.pmyhr8 Accessed from R via rgbif (https://github.com/ropensci/rgbif) on 2023-11-03 |
| *Thalictrum fendleri* | GBIF Occurrence Download https://doi.org/10.15468/dl.uj96tx Accessed from R via rgbif (https://github.com/ropensci/rgbif) on 2023-11-03 |
| *Trifolium longipes* | GBIF Occurrence Download https://doi.org/10.15468/dl.t5eps9 Accessed from R via rgbif (https://github.com/ropensci/rgbif) on 2023-11-03 |
| *Valeriana occidentalis* | GBIF Occurrence Download https://doi.org/10.15468/dl.mkz438 Accessed from R via rgbif (https://github.com/ropensci/rgbif) on 2023-11-03 |
| *Veratrum tenuipetalum* | GBIF Occurrence Download https://doi.org/10.15468/dl.8nnyf8 Accessed from R via rgbif (https://github.com/ropensci/rgbif) on 2023-11-03 |
| *Vicia americana* | GBIF Occurrence Download https://doi.org/10.15468/dl.cafyfn Accessed from R via rgbif (https://github.com/ropensci/rgbif) on 2023-11-03 |
| *Viola nuttallii* | GBIF Occurrence Download https://doi.org/10.15468/dl.up2xr5 Accessed from R via rgbif (https://github.com/ropensci/rgbif) on 2023-11-05 |
