## Appendix S2 for "Rare species do not disproportionately contribute to phylogenetic diversity in a subalpine plant community"

**Table S2:** Species in our research sites listed with replacement species used from Smith & Brown's ALLMB.tre phylogeny (see also Methods). Road is the lowest elevation site (2815 m), Pfeiler is the middle (3165 m), and PBM is the highest (3380 m).

| **RMBL species** | **Replacement from Smith & Brown 2018** | **RMBL site(s)** | **Data year** |
|---|---|---|---|
| *Senecio crassulus* | deleted | Pfeiler, PBM | 2022, 2021 |
| *Senecio integerrimus* | *Senecio triangularis* | Road, PBM | 2022 |
| *Cymopterus lemmonii* | *Cymopterus planosus* | PBM | 2022 |
| *Descurainia spp* | *Descurainia incana* | PBM | 2021 |
| *Potentilla gracilis* | *Potentilla gracilis var. flabelliformis* | Pfeiler, PBM, Road | 2022, 2021 |
| *Veratrum tenuipetalum* | *Veratrum virginicum* | Road, PBM | 2021 |
| *Agoseris glauca* | *Agoseris glauca var. dasycephala* | Pfeiler | 2021 |
| *Eucephalus engelmannii* | *Eucephalus breweri* | Pfeiler | 2021 |
| *Helianthella quinquenervis* | *Helianthella uniflora* | Pfeiler | 2022, 2021 |
| *Hydrophyllum capitatum* | *Hydrophyllum capitatum var. capitatum* | PBM, Pfeiler | 2022, 2021 |
| *Gayophytum diffusum* | *Gayophytum diffusum subsp. diffusum* | Road | 2021 |
| *Madia glomerata* | *Madia sativa* | Road | 2021 |
| *Primula pauciflora* | *Primula pauciflora var. pauciflora* | Road | 2022 |
| *Erigeron elatior* | *Erigeron grandiflorus* | PBM, Pfeiler | 2021, 2022 |
| *Penstemon crandallii* | *Penstemon crandallii subsp. crandallii* | Road | 2022 |
