## Appendix S3 for "Rare species do not disproportionately contribute to phylogenetic diversity in a subalpine plant community"

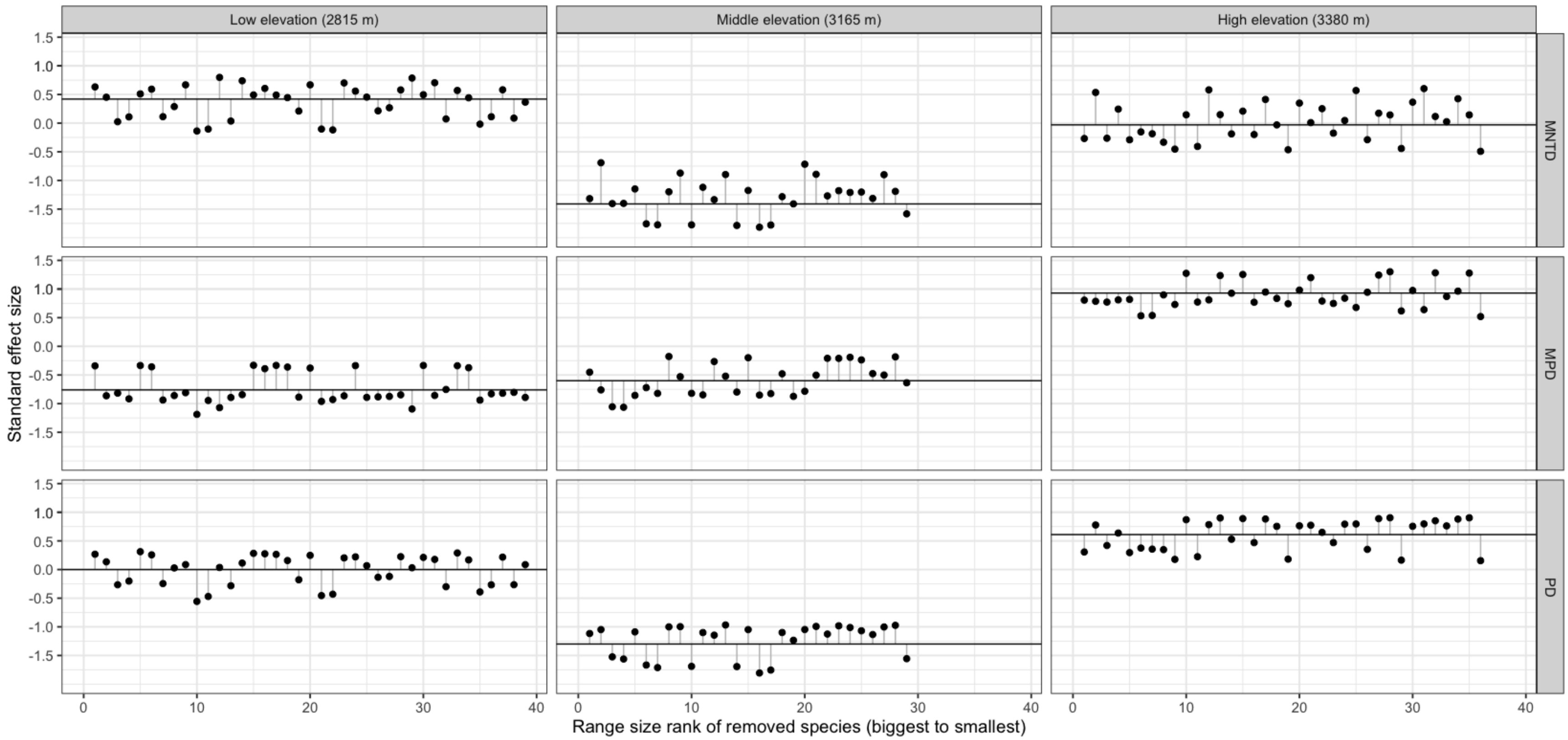


**Figure. S3:** Change in SES values when individual species are removed in order of decreasing range size. The horizontal black line in each figure shows the standard effect size of the metric for the overall community at each site, and dots show how the SES changes when the species at the rank indicated on the X axis is removed. MNTD values are shown in the top row, MPD values are in the middle row, and PD are on the bottom. The left column shows the low elevation site, middle column is the middle elevation site, and right is the high elevation site. There is no clear pattern suggesting that removing rarer species (right side of each panel) decreases MNTD, MPD or PD more than removing more common species (left side of each panel).
