## Appendix S4 for "Rare species do not disproportionately contribute to phylogenetic diversity in a subalpine plant community"

**Table S4:** P-values for autocorrelation tests at each site for both abundance and range size, all three phylogenetic metrics and 1, 2 and 3 steps.

| Site | Elevation | Categorization | Phylogenetic metric | Durbin-Watson statistic | Autocorrelation P | Lag |
|---|---|---|---|---|---|---|
| PBM | 3380 m | Abundance | MNTD | 2.578367 | 0.13 | 1 |
| PBM | 3380 m | Abundance | MNTD | 1.697201 | 0.326 | 2 |
| PBM | 3380 m | Abundance | MNTD | 2.082445 | 0.666 | 3 |
| PBM | 3380 m | Abundance | MPD | 1.891162 | 0.602 | 1 |
| PBM | 3380 m | Abundance | MPD | 2.518014 | 0.112 | 2 |
| PBM | 3380 m | Abundance | MPD | 2.092688 | 0.644 | 3 |
| PBM | 3380 m | Abundance | PD | 2.110638 | 0.852 | 1 |
| PBM | 3380 m | Abundance | PD | 1.917727 | 0.774 | 2 |
| PBM | 3380 m | Abundance | PD | 2.280713 | 0.29 | 3 |
| Pfeiler | 3165 m | Abundance | MNTD | 1.824263 | 0.47 | 1 |
| Pfeiler | 3165 m | Abundance | MNTD | 2.144139 | 0.632 | 2 |
| Pfeiler | 3165 m | Abundance | MNTD | 2.446184 | 0.172 | 3 |
| Pfeiler | 3165 m | Abundance | MPD | 2.04566 | 0.928 | 1 |
| Pfeiler | 3165 m | Abundance | MPD | 2.172034 | 0.624 | 2 |
| Pfeiler | 3165 m | Abundance | MPD | 1.922882 | 0.994 | 3 |
| Pfeiler | 3165 m | Abundance | PD | 2.078992 | 0.99 | 1 |
| Pfeiler | 3165 m | Abundance | PD | 2.192833 | 0.636 | 2 |
| Pfeiler | 3165 m | Abundance | PD | 2.445441 | 0.134 | 3 |
| Road | 2815 m | Abundance | MNTD | 2.575114 | 0.074 | 1 |
| Road | 2815 m | Abundance | MNTD | 1.651764 | 0.232 | 2 |
| Road | 2815 m | Abundance | MNTD | 1.71376 | 0.456 | 3 |
| Road | 2815 m | Abundance | MPD | 1.950843 | 0.728 | 1 |
| Road | 2815 m | Abundance | MPD | 1.906378 | 0.786 | 2 |
| Road | 2815 m | Abundance | MPD | 1.965041 | 0.962 | 3 |
| Road | 2815 m | Abundance | PD | 2.231123 | 0.588 | 1 |
| Road | 2815 m | Abundance | PD | 1.973191 | 0.892 | 2 |

| | | | | | | |
|---|---|---|---|---|---|---|
| Road | 2815 m | Abundance | PD | 1.60153 | 0.296 | 3 |
| PBM | 3380 m | Range size | MNTD | 2.475278 | 0.206 | 1 |
| PBM | 3380 m | Range size | MNTD | 1.885724 | 0.776 | 2 |
| PBM | 3380 m | Range size | MNTD | 1.449014 | 0.146 | 3 |
| PBM | 3380 m | Range size | MPD | 2.239795 | 0.576 | 1 |
| PBM | 3380 m | Range size | MPD | 1.792886 | 0.542 | 2 |
| PBM | 3380 m | Range size | MPD | 2.00366 | 0.836 | 3 |
| PBM | 3380 m | Range size | PD | 2.399343 | 0.304 | 1 |
| PBM | 3380 m | Range size | PD | 1.876702 | 0.716 | 2 |
| PBM | 3380 m | Range size | PD | 1.361636 | 0.062 | 3 |
| Pfeiler | 3165 m | Range size | MNTD | 2.082163 | 0.996 | 1 |
| Pfeiler | 3165 m | Range size | MNTD | 1.884824 | 0.764 | 2 |
| Pfeiler | 3165 m | Range size | MNTD | 1.806387 | 0.774 | 3 |
| Pfeiler | 3165 m | Range size | MPD | 1.755365 | 0.408 | 1 |
| Pfeiler | 3165 m | Range size | MPD | 2.267995 | 0.524 | 2 |
| Pfeiler | 3165 m | Range size | MPD | 1.637946 | 0.432 | 3 |
| Pfeiler | 3165 m | Range size | PD | 1.950825 | 0.764 | 1 |
| Pfeiler | 3165 m | Range size | PD | 2.125775 | 0.708 | 2 |
| Pfeiler | 3165 m | Range size | PD | 1.409608 | 0.162 | 3 |
| Road | 2815 m | Range size | MNTD | 2.153873 | 0.8 | 1 |
| Road | 2815 m | Range size | MNTD | 2.30677 | 0.364 | 2 |
| Road | 2815 m | Range size | MNTD | 1.625314 | 0.326 | 3 |
| Road | 2815 m | Range size | MPD | 1.491877 | 0.092 | 1 |
| Road | 2815 m | Range size | MPD | 1.846292 | 0.602 | 2 |
| Road | 2815 m | Range size | MPD | 1.930862 | 0.968 | 3 |
| Road | 2815 m | Range size | PD | 1.794683 | 0.386 | 1 |
| Road | 2815 m | Range size | PD | 2.453343 | 0.184 | 2 |
| Road | 2815 m | Range size | PD | 1.606169 | 0.3 | 3 |
