## Appendix S5 for "Rare species do not disproportionately contribute to phylogenetic diversity in a subalpine plant community"

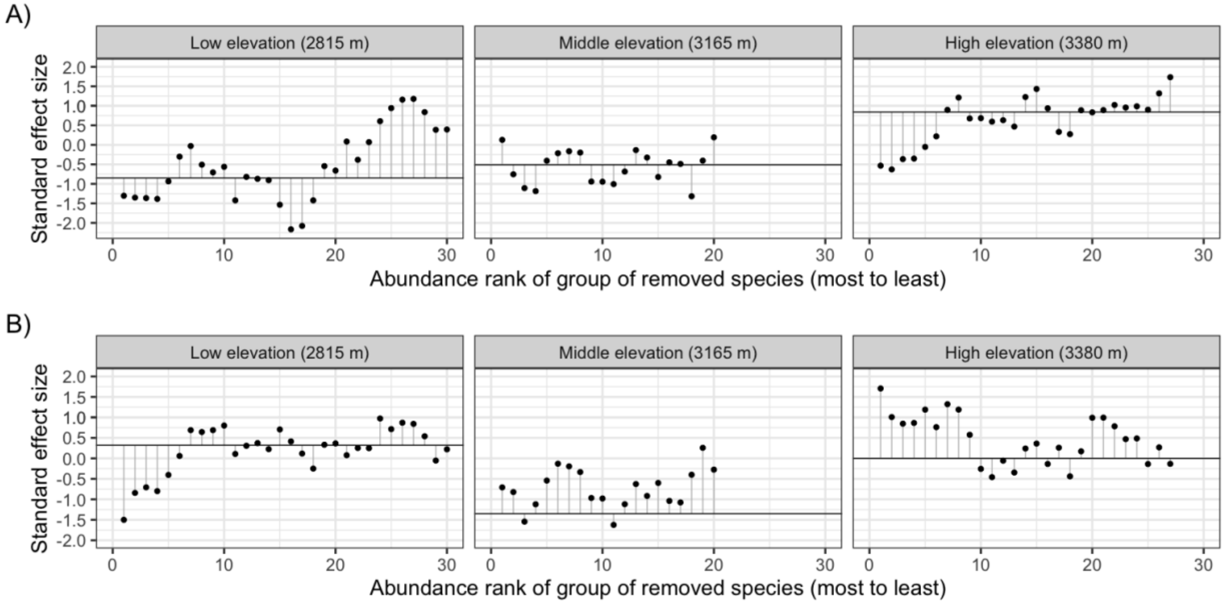


Figure S5: Change in SES values for MPD (A) and MNTD (B) when groups of ten species are removed in order of decreasing abundance. The horizontal black line in each figure shows the standard effect size of the metric for the overall community at each site, and dots show how the SES changes when the group is removed. Numbers on the x-axis are the ranking of the most abundant species in the group that was removed. For example, for the point at x=5, the species that were removed were ranked 5-14. The left panel shows the low elevation site, middle panel is the middle elevation site, and right is the high elevation site. There is no clear pattern suggesting that removing rarer species (right side of each panel) decreases MPD or MNTD more than removing more abundant species (left side of each panel).
